# Computational correction of cell-specific gene-independent effects in CRISPR-Cas9 essentiality screens: REStricted CUbic SplinEs with Mixed Models (RESCUE-MM)

**DOI:** 10.1101/2021.10.22.465496

**Authors:** Julie A. Dias, Shibing Deng, Vinicius Bonato

## Abstract

Increased gene copy number has been associated with a greater antiproliferative response upon genome editing, regardless of the true essentiality profile of the targeted gene. Many methods have been developed to adjust for genomic copy number technical artifacts. Existing methods use a two-step correction by pre-processing the data prior to the final analysis. It has been shown that two-step corrections can produce unreliable results, due to the creation of a correlation structure in the corrected data. If this structure is unaccounted for, gene-essentiality levels can be inflated or underestimated, affecting the False Discovery Rate (FDR). We propose a one-step correction using restricted cubic splines (RCS) to be a simpler alternative which reduces the bias in downstream analyses. Moreover, most existing methods combine guide-level results to yield gene-level estimates which can misrepresent the true gene essentiality profile depending on the guide-averaging method. Our model-based approach (RESCUE-MM) for copy number correction provides a more flexible framework that allows for guide-level essentiality estimation while accommodating more complex designs with grouped data. We provide comparisons to existing copy number correction methods and suggest how to include copy number adjustment in a one-step correction fashion in multiple experimental designs.

## Introduction

CRISPR (Clustered Regularly Interspaced Short Palindromic Repeats)-Cas9 screens allow for a systematic analysis of gene function. These screens are powerful tools in oncology, as they allow to establish gene essentiality profiles in the context of synthetic lethality and can also identify genes causing drug resistance. In order to evaluate gene essentiality profiles, genome-scale CRISPR-Cas9 knockout (KO) screens are performed in human cell lines. For such an experiment, an array of single-guide RNAs (sgRNAs) is designed, synthesized and then cloned into a lentivirus library. This library is later infected into the cancer cells at a low multiplicity of infection (MOI) to ensure only one sgRNA copy is integrated per cell. Each sgRNA is made up of approximately 20 nucleotides and is complementary to its target region guiding the Cas9 protein to its desired cutting point. The Cas9 protein then makes a double-strand DNA break (DSB) which often results in the knockout of the targeted gene [1]. The cells are then left to grow under various experimental settings. After cell selection, the read-count level of each sgRNA is quantified by next generation sequencing to determine relatively depleted and enriched sgRNAs.

It has been established that knocking out genes in copy number amplified genomic regions tends to generate the greatest disruption in cell growth, regardless of the gene’s true biological essentiality (**Fig 1A**). Indeed, inducing a DSB in a copy number amplified region is likely to trigger DNA damage response mechanisms and cause cell-cycle arrest [2], presumably, in addition to the targeted gene’s fitness role, as seen in focal copy number aberrations, for example [3]. This is illustrated by a high gene essentiality score which in turn increases the number of false positives in CRISPR-Cas9 KO screens [4], especially in cancer cell lines where large copy number alterations are common. Several methods have been developed to account for gene-independent copy number related artifacts on sgRNA depletion/enrichment counts, using piecewise linear splines [5], generalized additive models [6] or univariate linear models [7]. While the true relationship between the sgRNA depletion and copy number remains unknown, most suggest it is non-linear and cell-specific [8].

**Fig 1.**
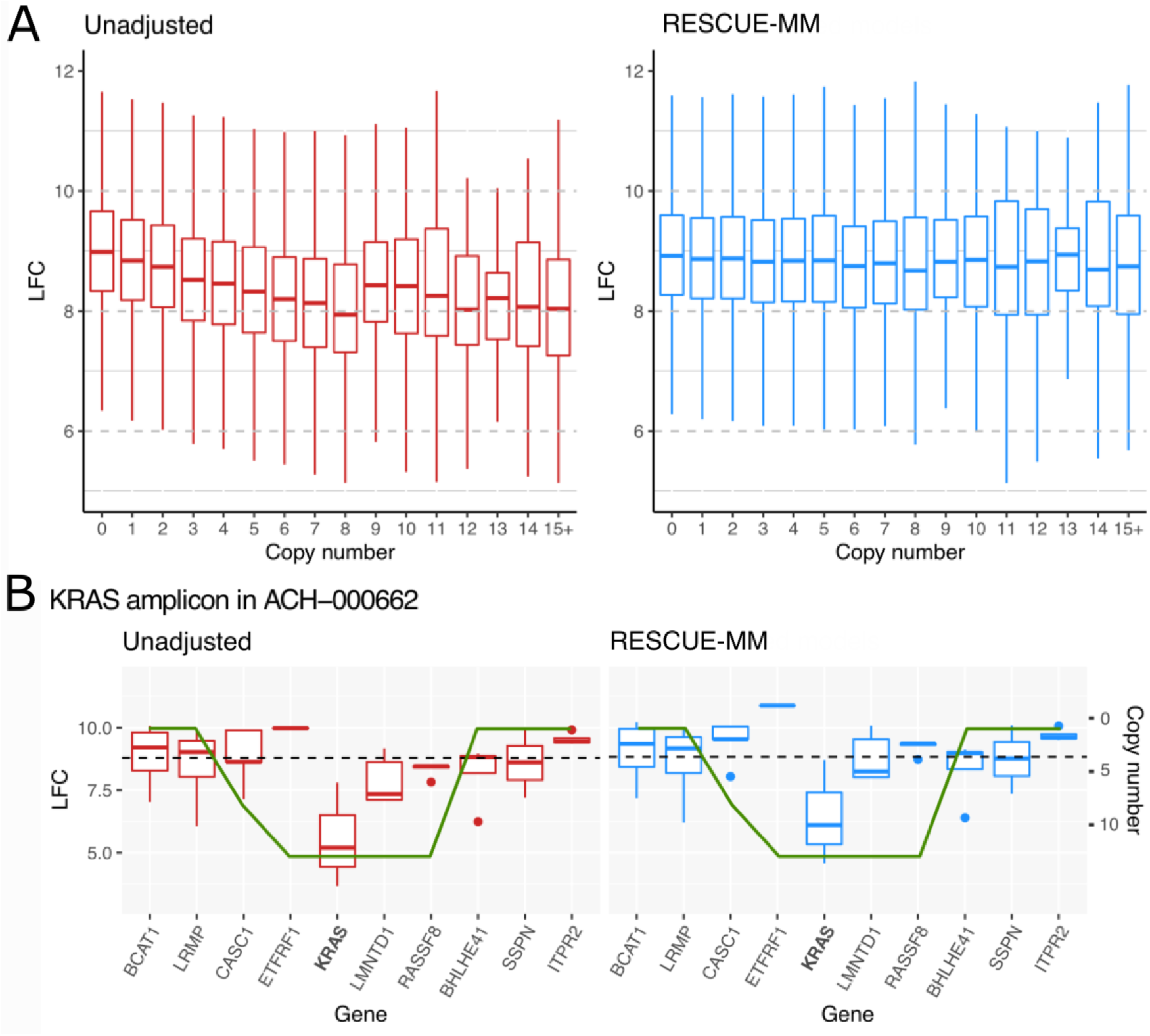
Copy number effect on CRISPR-Cas9 knockout sensitivity and RESCUE-MM correction. A) LFC across 112 cancer cell lines summarized by a boxplot for each rounded copy number value. B) LFC within the KRAS amplicon in the ACH-000662 lung cancer cell line summarized by a boxplot for each gene in the amplicon. The green line displays the copy number value scaled to the data. The left panel displays the unadjusted LFC, while the right panel shows the LFC after RESCUE-MM correction. The black dashed line represents the mean LFC across all guides and cell lines for the unadjusted LFC (left) and RESCUE-MM corrected LFC (right).

All of these existing methods rely on a two-step procedure to first adjust for copy number effect and only then investigate synthetic lethality associations with oncogenes. This two-step procedure can often result in an exaggeration or reduction of statistical significance, symbolized by an increased FDR or an inability to recover known synthetically lethal genes. This is due to the creation of a correlation structure when using all data points within a genomic region [9]. Our method is a one-step procedure which can test for associations of sgRNA depletion/enrichment and common oncogenic drivers while simultaneously adjusting for the non-linear gene-independent copy number effects using restricted cubic splines (RCS) [10]. Our approach, REStricted CUbic SplinEs with Mixed Models (RESCUE-MM), although simple, is very flexible to accommodate modelling of data originated from complex designs. For example, data sets with multiple cell lines and multiple replicates can be modelled in a one-step approach while allowing each cell line to have its own copy number sensitivity in highly amplified genomic regions as well as accounting for replicate variability. Indeed, systematically averaging over replicates may not be appropriate in the case of heterogeneous within-sample variability [11]. Moreover, RCS are also flexible enough to be used in the generalized linear model framework and, therefore, count data can be modelled by a negative binomial model (RESCUE-GLMM) [12]. Finally, some methods have a strict weighting method to average guide scores yielding an overall gene-level essentiality that often discards low efficacy guides [5], while our method allows for an arbitrary weighting of the guide-level scores.

## Results

### Adjusting for copy number

As expected, RESCUE-MM notably reduces the relationship between copy number and log-fold change (LFC) found in the unadjusted guide LFC (**Fig 1A**). Concentrating on NSCLC focal copy number aberrations, ie, oncogenes whose amplification is associated to the progression of cancer, such as KRAS, RESCUE-MM moderately reduces the relationship between counts and copy number but preserves the depletion in counts when the gene is located in a genomic amplified region (**Fig 2A**). Indeed, it has been found that KRAS is frequently amplified, and that this amplification is correlated with activating mutations of KRAS [13]. Thus, count depletion should only be markedly present in amplified oncogenes. In contrast, RESCUE-MM is able to markedly remove the copy number effect in non-essential amplified genes such as ASZ1 (**Fig 2B**).

**Fig 2.**
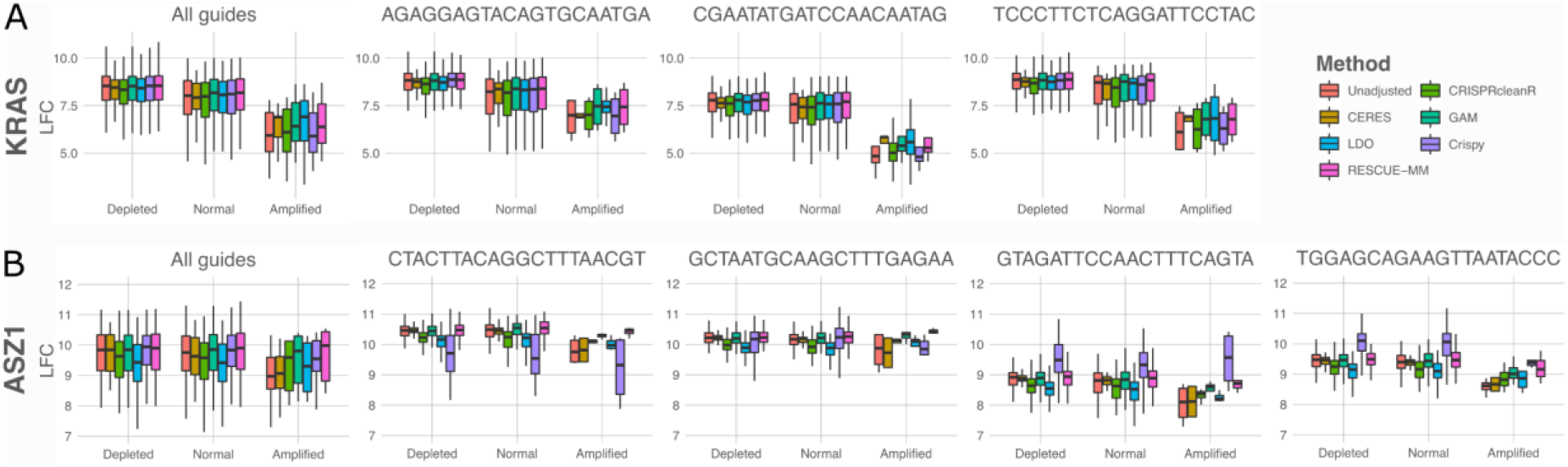
Copy number effect comparison according to gene category at a gene-level and guide-level, according to copy number category (Depleted < 1.2, 1.2 ≤ Normal < 5, Amplified ≥ 5). A) KRAS (essential oncogene amplified in lung cancer) LFC depletion comparison across 112 cancer cell lines. B) ASZ1 (non-essential gene with high copy number) LFC depletion comparison across 112 cancer cell lines.

To further investigate the performance of our approach in well-characterized NSCLC amplicons, we focused on a KRAS amplicon where KRAS has a copy number of 13. In **Fig 1B**, the LFC before and after RESCUE-MM correction are shown for the KRAS amplicon in ACH-000662. In **Fig 1B** (left), CASC1, LMNTD1 and RASSF8 show equivalent LFC levels, assumed to be caused by the copy number artifact. In contrast, KRAS shows a markedly lower LFC. Upon RESCUE-MM correction, shown in **Fig 1B** (right), the copy number effect has been correctly removed for CASC1, LMNTD1 and RASSF8 but the KRAS LFC remains low. This identifies KRAS as the amplicon driver, as expected [14]. Moreover, RESCUE-MM decreases the correlation between neighbouring genes, allowing for a better detection of synthetically lethal genes and reducing the false discovery rate (**Suppl. Table 1**). Methods like CERES markedly increase the correlation between neighbouring genes, which can produce unreliable corrected counts, especially in copy number amplified regions.

### Between and within cell line variability

Methods such as CERES, CRISPRcleanR, Crispy, LDO and GAM use various approaches to adjust for copy-number (see Methods). We hypothesize that the best methods will preserve the between and within cell lines variability for control guides and decrease the between-cell line variability while preserving the within-cell line variability for guides in copy number amplified genomic regions. The portion of the between-cell line variability for guides targeting copy number amplified genomic regions can be explained by the copy number off-target effects and should generally be successfully removed after copy number adjustment. In order to compare the performance of RESCUE-MM and other existing copy number adjusting methods we selected guides that show higher overall variability in the unadjusted data as well as a copy number effect across cell lines.

When applying RESCUE-MM, the coefficient of variation (%CV) between and within cell line for control guides remains unchanged between the unadjusted and copy number adjusted log-counts. However, for guides potentially affected by copy number artifacts, the %CV is successfully reduced between cell lines after the adjustment, especially on the lower %CV range. Moreover, the %CV for within cell line variability is preserved in these potential copy number affected guides **(Fig 3)**. In comparison to the RESCUE-MM method in control guides, the %CV within and between cell lines is similar or higher when using LDO, GAM, Crispy and CRISPRcleanR **(Suppl. Fig 1)**. Finally, the total variability is higher in LDO, GAM and Crispy. The within cell line %CV when using Crispy is marginally better preserved but a large discrepancy between methods is observed. With CRISPRcleanR, the total variability appears relatively comparable except for guides with relatively higher %CV **(Suppl. Fig 1)**.

**Fig 3.**
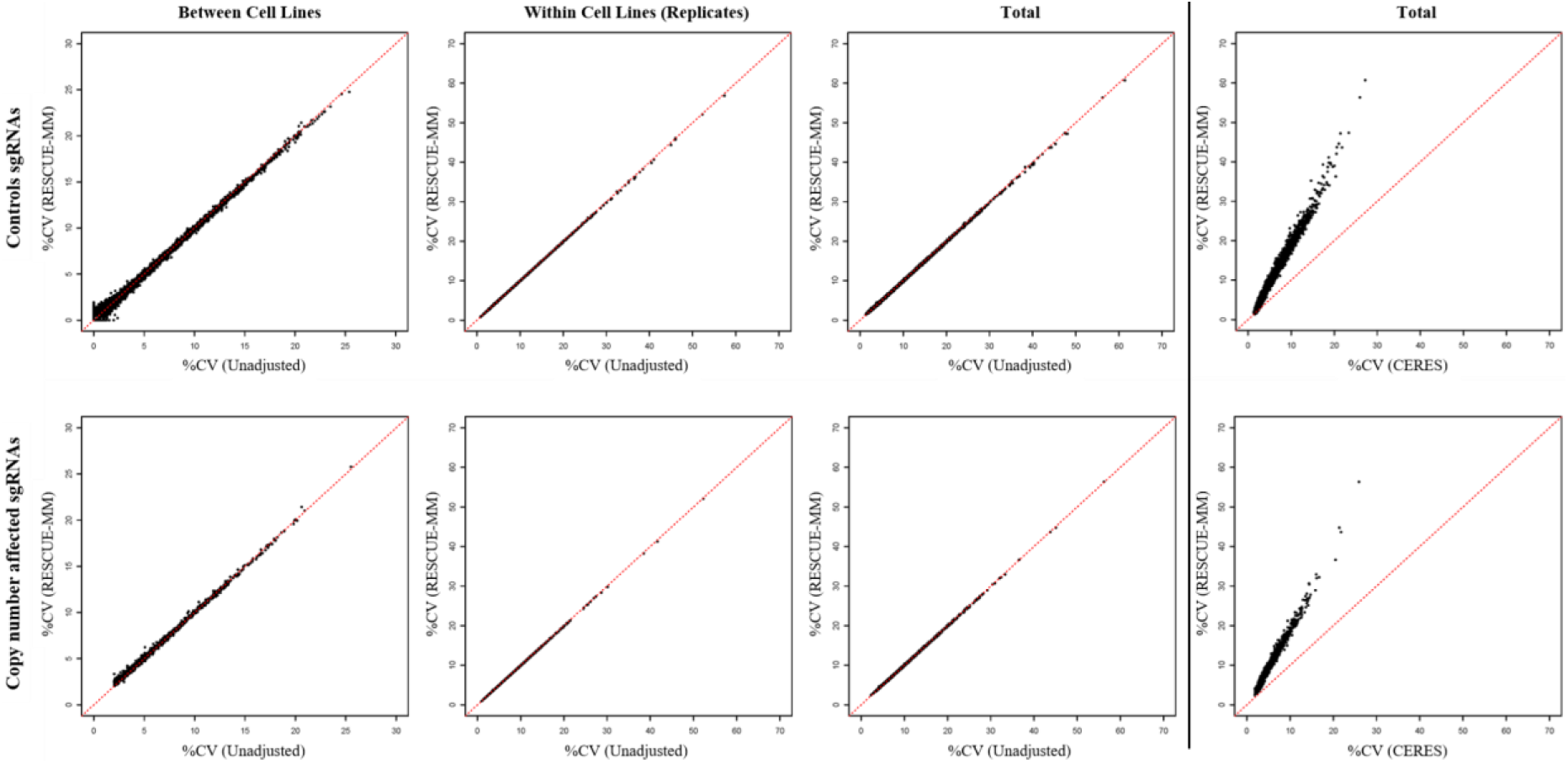
Comparison in the variance of LFC following copy-number adjustments. From left to right: between cell lines variation comparison, within cell lines variation (i.e. within replicates), total variation (within variation + between variation). The first three columns from the left compare unadjusted counts and RESCUE-MM corrected counts while the rightmost column compares CERES counts and RESCUE-MM corrected counts. Each point represents a guide and for each guide the %CV is computed. Top row compares %CV in control guides, bottom row compares %CV in copy number affected guides.

For guides potentially affected by copy number artifacts, the %CV within cell lines is higher in GAM and Crispy. LDO appears to perform similarly, but some of the guides show some slight discrepancies. Moreover, in comparison to CRISPRcleanR, RESCUE-MM seems to reduce between cell line variability in the lower range of %CV shown although this may still be inconclusive since some of the guides show some moderate discrepancies **(Suppl. Fig 2)**. CERES outperforms RESCUE-MM in total variability in control and copy number affected guides, but this may be due to this method artificially reducing variability by averaging samples and lacking flexibility in its copy number modelling approach which may force a fit where there is no copy number effect thus flattening differences across cell lines regardless of the copy number influence **(Fig 3)**.

Overall, RESCUE-MM seems to be a better fit for the copy number effect and outperforms most existing methods. In some instances, other methods appear to perform better due to a rigid copy number fit. The large reduction in total variability when applying CERES is in part due to its intentional lack of replicate variability which can artificially reduce the %CV. Methods like Crispy and CRISPRcleanR systematically force a copy number effect, even in the absence of true copy number effects. This may be imposing an exaggerated similarity between replicate guides within samples in controls.

### Association with mutation profile data

Residuals after correction for copy number were gathered and tested for association with genetic alterations (mutations/deletions/amplifications) across 89 of the 112 lung cancer cell lines. Mirroring the other existing copy number correcting methods, a single-step RESCUE-MM copy-number correction was used. On a quantile–quantile plot (QQ-plot) of the observed −log10(P) from different methods against the expected −log10(P) under the null condition, control guides are expected to follow the identity line. Focusing on the association of control guides and CDKN2A deletion (the most frequent alteration in the cell line subset), significance is notably inflated by CRISPRcleanR and deflated by CERES. Reassuringly, our method follows the unadjusted count distribution, showing control guides are not significantly associated with genetic alterations (**Fig 4A**).

**Fig 4.**
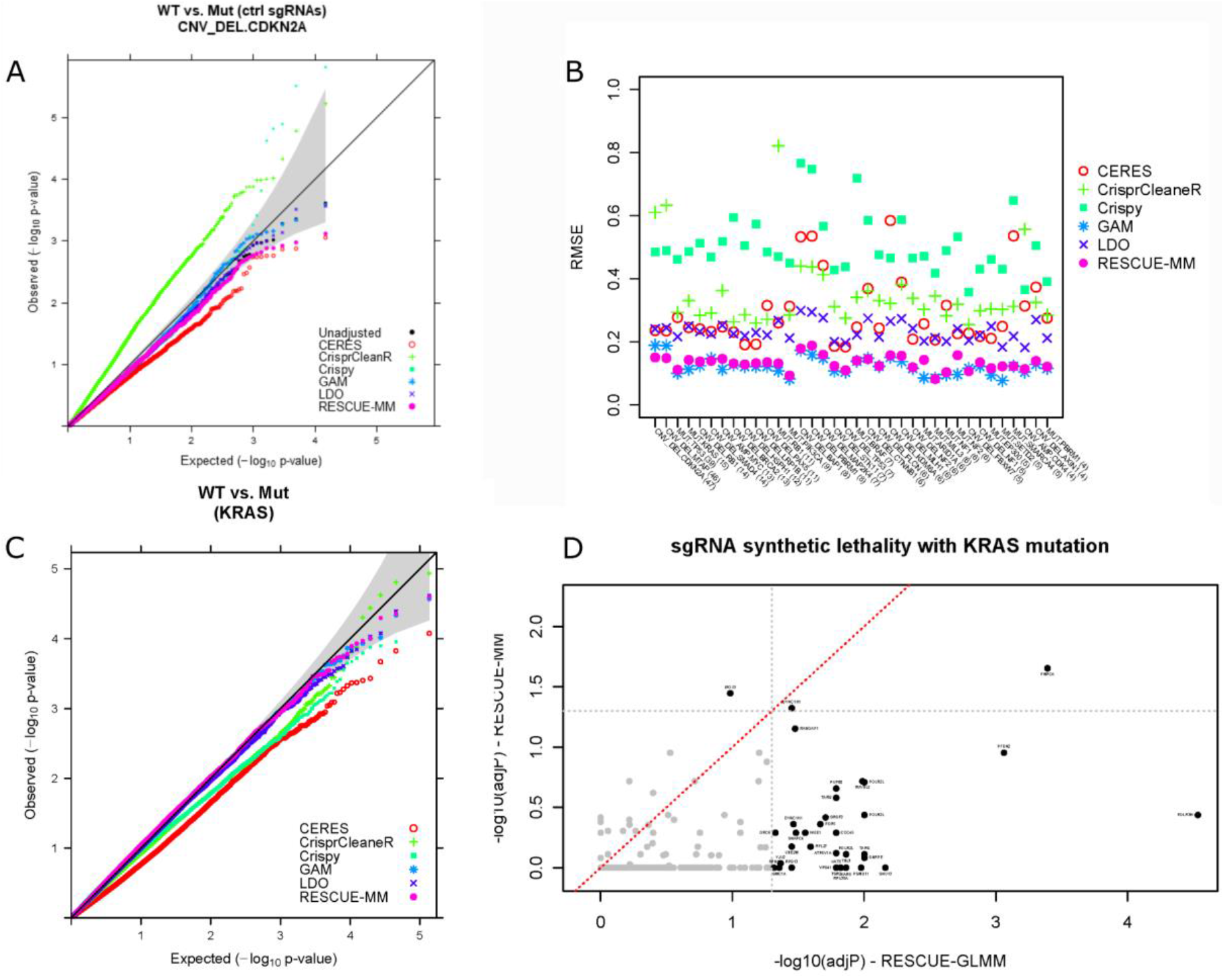
Comparison of p-value inflation/deflation testing association between adjusted LFC and genomic alterations (deletion, amplification, mutation) A) QQ-plot comparing significance of corrected LFC and a deletion in CDKN2A in control guides. B) Root mean square error (RMSE) between unadjusted LFC and adjusted LFC for existing copy number correcting methods and RCS in control guides. C) QQ-plot comparing significance of corrected LFC and a mutation in KRAS in all guides using 2-step approach methods. D) Scatter-plot comparing Benjamini-Hochberg adjusted significance of corrected LFC and a mutation in KRAS in all guides using one-step approaches. Dashed grey lines show the cut-off for significance.

In **Fig 4B**, the RMSE between significance of the unadjusted counts p-value and the copy number corrected counts is shown for each copy number correcting method for the top 36 genomic alterations, representing the magnitude of p-value inflation/deflation in control guides. In the same fashion, RESCUE-MM model fitting mirrors the other existing copy number correcting methods, using the residuals after adjustment as inputs for the association tests (two-step approach). With high RMSE, methods like Crispy, CERES, and CRISPRcleanR systematically inflate/deflate the significance of genetic alterations and copy number corrected counts in control guides. On the other hand, RESCUE-MM performs appreciably better with consistent low RMSE, similarly to GAM.

### Flexibility to be fully model based

As our data show **(Fig 4B)**, applying copy number correcting methods on CRISPR-Cas9 gene knockout data can produce unreliable results (exaggerated significance) due to using a two-step rather than joint inference procedure. This is particularly relevant when the genomic copy number of the cell lines are unevenly distributed, which is the case for some genes more likely to be amplified (particularly in cancer cell lines). The magnitude of the significance exaggeration or understatement depends on how unbalanced the gene (or guide)-copy number design is, rather than on the sample size in itself. If copy number differences are not constant across all genes and/or guides, this problem will persist no matter how large the sample size. This increases the number of false-positives in copy number-amplified regions when analysing the normalized data: removing the copy number bias in two steps introduces a correlation structure in the data as the copy number mean and/or variance are usually estimated using all the data corresponding to the guides mapping to the particular gene, and that across cell lines. Thus, all adjusted data points within a guide and even within a gene are correlated with each other. This correlation structure is responsible for the increased (or sometimes decreased) significance in the results. A one-step procedure has the advantage of including gene copy number effects directly in the modelling and analysis step, and using a mixed model allows to remove this effect independently for each cell line.

Replicate measurements can increase statistical power depending on the number of replicates and the correlation across replicates [15]. However, some existing copy-number adjusting methods such as CERES average replicates before correlations are calculated. Such practice is not recommended if there is poor concordance between cell lines and the within-sample variability is heterogeneous [16]. Indeed, averaging over replicates should only be considered if the within cell line correlation is very high.

KRAS mutations are known to be oncogenic in humans, and to play a role in the growth of cancer cells, particularly in lung cancer [17]. Two-step approaches are not able to identify any significant hit after multiple hypotheses testing adjustment (**Fig 4C**) whereas, as shown in **Fig 4D**, linear and generalized linear model versions of RESCUE algorithm (respectively RESCUE-MM and RESCUE-GLMM) were able to identify multiple significant hits (using the Benjamini-Hochberg multiple hypotheses testing adjustment) when testing for association. Among many hits representing synthetic lethality with a KRAS mutation we found the DYNC1H1 gene which is intricately involved in moving cell materials along microtubules and has previously been identified as a driver for progression and metastasis of colorectal cancers [18]. Likewise, we identify RANGAP1, a member of the Ras superfamily which regulates nucleo-cytoplasmic transport of molecules, which disruption has been linked to cancer proliferative signalling, resistance to apoptosis, and invasion/metastasis [19]; IPO13, a mediator of docking of the importin/substrate complex to the nuclear pore complex by a Ran-dependent mechanism, which inhibition is thought to be a potential therapeutic strategy for NSCLC [20]; and, RUVBL2, a helicase belonging to the INO80 multi-subunit protein complex, which is found to have tumour-promoting roles in several cancers and has been associated with opening the chromatin state in cancer cells and enhancing oncogenic transcription [21]. In addition, we were able to identify 28 other genes whose biology allows for potential synthetically lethality in the presence of a KRAS (**Suppl. Table 2**).

## Methods

### Quality control and pre-processing of CRISPR-Cas9 samples

Raw read counts of CRISPR pooled library screens for 257 lung cancer cell lines samples (112 lung cancer cell lines, 1-4 replicates per cell line) were obtained from the Broad DepMap project data depository (Data release 20Q2 - https://depmap.org/portal/download/). For efficiency purposes, samples used in this manuscript were limited to lung cancer cell lines, the most represented group in DepMap. Cancer vulnerabilities were investigated in these cell lines using the genome-wide Avana library to knock out 18,524 genes using 72,133 sgRNAs. To avoid genetic interaction effects, sgRNAs targeting multiple matched chromosomal regions were filtered out from the analysis in this manuscript, resulting in a total of 68,119 sgRNAs targeting 17,951 genes (~3.8 sgRNAs per gene). Raw read counts were subsequently normalized using the median of ratios method of normalization [22]. A pseudo-count of one was added to all normalized counts prior to log2 transforming the data to meet the distributional assumptions of normality of the methods investigated in this manuscript. Additional pre-processing steps were undertaken to remove undesired variation in the data attributable to experimental artifacts. Technical artifacts as processing batches (3 batches), intrinsic cell line Cas9 activity (categorized in deciles), cell line null-normalized mean differences categorized in deciles (see [23] for definition), and culture types (adherent, suspension, or mixed) were corrected for in the dataset at the sgRNA count level using the ComBat method [24] available in the *sva* R package [25]. These log2 transformed artifact adjusted counts represent the post library-transfection quantities for individual sgRNAs (LFC). Departures from central values may indicate relative sgRNA depletions or enrichments.

### Copy number estimates and genetic alteration profiles

Genome-wide copy number data for the matching 112 lung cancer cell lines were downloaded from the DepMap data portal (https://depmap.org/portal/download/). Copy number log2 transformed estimates, given by chromosomal segments, were matched to sgRNAs in each cell line. In addition, gene-level alterations for 89 of these cell lines were downloaded from the COSMIC database (https://cancer.sanger.ac.uk/cosmic/download) and the 36 most common ones were tested for associations with individual sgRNAs depletions/enrichments. Gene-level genomic alteration status was summarized per cell line as: mutation - presence (1) vs. absence (0) of any functional mutation in the genomic span of a gene of interest; deletion - presence vs. absence of any genomic region with copy number ≤ 1.2; or amplification - presence vs. absence of any genomic region with copy number > 6. In total, 13 driver mutations, 21 chromosomal deletions, and 2 focal copy number amplifications were tested for synthetic lethality in this study (**Fig 4B**).

### Adjusting for copy number using restricted cubic splines

The existence of a relationship between sgRNA counts depletion and copy number alterations has been widely established and a vast array of modelling methods from piecewise linear splines [5] to generalized additive models [6] have been employed to adjust the copy number effect and, consequently, reduce false discoveries in these copy number amplified regions. Alternatively, modelling the relationship between LFC and copy number as non-linear and cell line specific using restricted cubic splines [10] is proposed and evaluated in this manuscript. Cubic polynomials have been found to have good ability to fit sharply non-linear relationships and, in comparison to other copy number adjustment approaches, are thought to be more flexible to be explored in additional modelling contexts [10]. A cubic spline can be viewed as a piecewise cubic polynomial, where the number of “pieces” is dictated by the number of knots *k* used. Within each piece is a cubic polynomial, with restrictions to ensure the spline is continuous and smooth at each knot. Restricted cubic splines constrain the function to be linear before the first knot and after the last knot to avoid poor fitting in the tails [10][26]. In the linear model context, the RCS structure with *k* knots will have *k* components: one constant value (the intercept, *β*_0_), one component (*β*_1_) which is linear in the copy number variable, and *k* – 2 non-linear (cubic) components (*β_m_*) in the copy number variable:

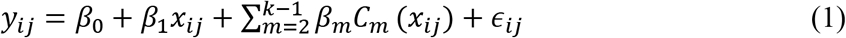

where *y_ij_* is the known vector of LFC values for sgRNA *i* in cell line *j*, *X_ij_* is the log2 copy number at the genomic location of sgRNA *i* in cell line *j*, and *∈_ij_* is the residual error which is normally distributed and centered at zero (*i*=*1*,…, *68119*; *j*=*1*,…, *112*). In this equation, *C_m_* is the cubic component that falls in the *m*th window, where *m* ∈ {1, *k* + 1}. The rcspline.eval function available in the *r*ms (version 4.5-0) R package was used in default mode (k = 5 knots positioned at the following *x*-quantiles: 0.05, 0.275, 0.5, 0.725, and 0.95) to calculate the design matrix containing the 3(*k* – 2) non-linear spline variables denoted by *C_m_* in equation (1). The reader is referred to the texts of [10] and [27] for discussions on why this choice of number and position of knots is considered adequate for the sample size used in this study.

### Flexibility to model complex experimental designs

As an extension of model (1), the linear mixed modelling framework [11] flexibilizes the assumption of equal copy number effects (*β*) across cell lines by allowing now each cell line *j* to have its own vector of copy number effects (*b_j_*) as

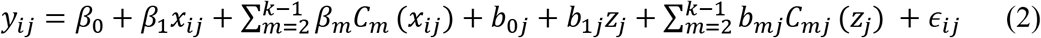

Here, the k*1 vector *b* is normally distributed, centered at zero and with variance-covariance matrix *ψ*. The square, symmetric, and positive semidefinite matrix with independent random effects *ψ* is represented as

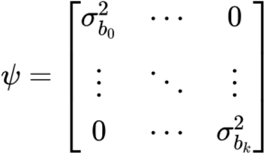

The adjusted LFC counts for each sgRNA *i* (for simplicity sgRNA index is omitted subsequently when redundant) can then be obtained by keeping the fixed effect intercept as well the residuals from the linear mixed-effect model, yielding

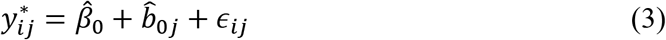

The lmer function in the *lme4* (v1.1-26) R package was used to estimate the model parameters with the restricted maximum likelihood algorithm.

### One-step *vs*. two-step approaches

For each genomic alteration, adjusted LFC values 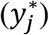 can subsequently be used for uncovering synthetic lethality hits with this oncogenic alteration in a two-step approach by testing the null hypotheses *H_0_:λ* = 0 for each sgRNA *i* as in the model below

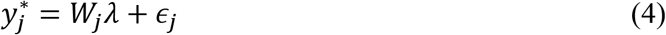

where, *W_j_* is the indicator variable for the presence of the genomic alteration (deletion, amplification, or mutation) in cell line *j* and *λ* describes the magnitude of the synthetic lethality effect between that genomic alteration and sgRNA *i*.

Alternatively, the synthetic lethality (fixed) effect for each genomic alteration can be modelled in a one-step approach (RESCUE-MM) directly from (2) with the addition of fixed effects for sgRNAs (*γ_i_*), genomic alteration (*Ʌ*), and interaction of sgRNA *i* and genomic alteration (*γ_i_Ʌ*), such as

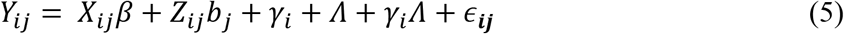

where *γ_i_* is the indicator for the sgRNA being tested and *Λ* is the indicator variable for the genomic alteration being tested. Similarly, the null hypotheses *H_0_*: *γ_i_Λ* = 0 are tested and p-values obtained by the one-step vs. two-step approaches were compared for identification of any biases.

### Flexibility to model sequence counts

RCS flexibility also allows the modelling of the non-linear relationship between copy number and the original sgRNA sequence counts directly, therefore preserving the original assay measurements and data distribution. From equation (5), it follows that the negative binomial generalized linear model (RESCUE-GLMM) can be written as

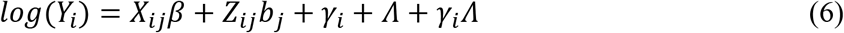

and estimates for *γ_i_Λ* were obtained in a one-step approach using the glmer function for the negative binomial family available in the *lme4* (v_1.1-26) R package. The p-values for *γ_i_Λ* generated by the linear (5) and generalized linear (6) models were compared.

### Comparison with existing copy number-adjusting methods

An exhaustive literature search identified the methods listed below as commonly used CN-correcting methods for CRISPR screen assays which were thus chosen to be compared in terms of performance and model flexibility to the RESCUE-MM alternative proposed here. All methods are explained in Table 1.

**Table 1:** Existing copy number-adjusting methods and their characteristics. Running time is based on 257 samples, 68,119 sgRNAs targeting 17,951 genes (MacBook Pro 2019 2.6GHz 5-Core Intel Core i7).

| Algorithm | CERES<br>(Meyers et al.) | CRISPRcleanR<br>(Iorio et al.) | LDO<br>(De Weck et al.) | GAM<br>(De Weck et al.) | Crispy<br>(Gonçalves et al.) |
| --- | --- | --- | --- | --- | --- |
| Published in | 2017 | 2018 | 2018 | 2018 | 2019 |
| Methodology | The measured individual sgRNA depletion is modelled as a sum of the gene-knockout effect and the copy-number effect. The copy-number effect is modelled by a piecewise linear spline, evaluated at the number of genomic sites targeted, determined by the target loci and the CN at each locus. For the gene-level knock-out effect, the cumulative depletion effects of all guides targeting the same gene are scaled by a guide activity score, restricted to values between 0 and 1, to capture and mitigate the influence of low-quality reagents. Replicate samples are averaged before being computationally modelled. | A circular binary segmentation algorithm is applied to the genome-wide patterns of sgRNA LFC across individual chromosomes in a cell line. It then detects genomic segments containing multiple sgRNAs with sufficiently equal LFC. If these segments contain sgRNAs targeting a minimum number of distinct genes then the sgRNA in the segment are most likely responding to CRISPR-Cas9 targeting in a gene-independent manner, and LFC values are corrected via mean-centering. | The phenotype scores for each guide are corrected by considering guide scores targeting the other genes in its direct genomic neighbourhood to assess each guide's true essentiality. Neighbouring guides showing no phenotypes or "neutral" gene guides are used to fit a regression tree to estimate the copy number effect. The copy number effect identified by the regression is then removed from the original essentiality score to reveal the true essentiality score. | The screening data is modelled as a function of the known copy number values and removes the systematic effect from the measured lethality. The model consists of smoothing functions estimated by non-parametric means from the data. | Gaussian processes regression is used to model the non-linear effects between the segment copy-number ratio (segment copy-number divided by the estimated chromosome copy-number) and the CRISPR-Cas9 fold changes. The squared-exponential kernel (RBF) is fitted independently for each sample to guarantee that CRISPR-Cas9 global effects that are sample specific are captured automatically by the defined kernel. |
| Running time | ~17 hours | ~70 mins | ~250 mins | ~4 mins | ~30 mins |
| Two step procedure | Yes | Yes | Yes | Yes | Yes |
| Requires copy number data | Yes | No | No | Yes | Yes |

Intra and inter cell line variability of 7393 sgRNA controls and 1520 sgRNAs showing large off-target copy number effects were selected to illustrate the comparisons of RESCUE-MM with the methods described above. sgRNA controls target essential and non-essential genes as listed in [5]. sgRNAs with large off-target copy number effects were identified as directly associated with copy number (p < 0.1) and highly variable (%CV > 2) across cell lines in the unadjusted data.

Additionally, sgRNA controls and sgRNAs with large off-target copy number effects were tested for synthetic lethality versus the top 36 most common genetic alterations in this set of cell lines. Unadjusted LFC values and adjusted LFC values obtained by each comparing method were univariately tested for association and inflation or deflation of p-values was investigated by comparing the p-values generated by each method to the ones obtained before any type of copy number effect adjustment.

## Discussion

High-throughput CRISPR-Cas9 screens are becoming paramount in the field of oncology and personalized medicine, with the potential to identify synthetically lethal genes and genes responsible for drug sensitivity/resistance. However, these screens are often affected by gene-independent antiproliferative effects such as copy number artifacts. As shown in previous studies, sgRNAs targeting genes that are amplified at high copy number are negatively correlated with cell proliferation, regardless of the gene’s true essentiality. In this report, we describe the restricted cubic splines (RCS) with mixed models method, namely RESCUE-MM, to correct for copy number bias, allowing to reduce the number of false positives and ease the identification of true positives, while allowing for model flexibility. The counts are corrected using a mixed-effect model where the copy number artifact is modelled using RCS. Compared to methods that model the relationship between the sgRNA depletion and copy number artifacts as linear piecewise, cubic splines are smoother and provide a better fit with the advantage of being robust at the tails where relatively large copy number gains or deletions could show instability. We applied our method (RESCUE-MM) to a subset of the Broad DepMap screening data, with a total of 18,524 genes across 112 lung cancer cell lines. RESCUE-MM correction markedly reduces the number of false positives by adjusting the depletion of counts caused by a copy number effect in non-essential genes, as was shown by the adjustment of ASZ1-targeting sgRNA counts in cell lines where ASZ1 is amplified. The ability of RESCUE-MM to preserve known cancer gene dependencies within and outside of amplified genomic segments was shown by the KRAS amplicon in ACH-000662 where the KRAS dependency was recovered, while removing the false positives of LMNTD1 and RASSF8. As RESCUE-MM’s copy number correction allows for each cell line to have its own copy number effect, we assume RESCUE-MM works efficiently regardless of the analysed dataset’s sample size, including single sample experiments. This fact indicates that RESCUE-MM could be implemented as an alternative for copy number adjustments in methods currently used for single cell line experiments such as MAGeCK which currently rely on copy-number adjustments performed after hypothesis testing [28].

Our motivation for developing a flexible copy-number adjustment method came from the observation that existing methods either artificially reduced guide-level variability by averaging replicates and/or lacked flexibility in their copy number modelling approach forcing a fit where there is no copy number effect leading to flattened differences across cell lines regardless of the true copy number influence. Moreover, most methods use a two-step procedure to infer gene essentiality profiles, by correcting for the copy-number effect in a separate step from testing for association with genomic alterations. Our method allows to increase the significance of true positive hits and reduce that of false positive hits, by having the flexibility of each cell line to have its unique copy number effect, while directly modelling counts as a function of copy number and mutation status in one step. Despite its practical benefits, log-transforming counts data to achieve normality can bring problems such as suboptimal variance stabilization which directly impact p-values [29]. In addition, our method allows to correct counts at a guide-level, giving the flexibility of weighing each guide’s count arbitrarily.

The significance of control guides should follow that of a uniform distribution and most methods including CERES, Crispy, and CRISPRcleanR systematically inflate or deflate p-values representing the association between genomic alterations and residuals after correction for copy number. RESCUE-MM, similarly to GAM, preserves the expected significance within control guides, showing it successfully reduces the number of false positives.

When testing for association between gene knockout and genomic alterations via a one-step procedure, modelling copy number as a random effect using RESCUE-MM and allowing for each cell line to have its own copy number effect, we saw a distinguishable improvement in significance. Taken individually, these improvements are incremental but important, especially in the case of multiple-hypothesis testing where p-values are adjusted (using Benjamini-Hochberg correction, for example). This one-step modelling approach allows for a better detection of true positives and reduces the number of false positives seen in other methods.

Given its performance for the Avana library, we believe RESCUE-MM performs equally in other genome-wide libraries such as the Brunello [30] and Whitehead [31] libraries. Our method requires copy number data for each cell line, unlike other methods like CRISPRcleanR. While methods which do not require orthogonal data may be advantageous when reliable copy number information is not available, having copy number information allows for a more informed fit when using smaller (gene or pathway specific) libraries. Furthermore RESCUE-MM is easily implemented and runs rather quickly. Other methods like CERES which use complex modelling schemes result in lengthy running times, approximately 40 times as long as RESCUE-MM. We suggest that complex and/or slow running models do not add benefit to the final analysis and that simpler and faster methods should be preferred.

## Acknowledgments

We thank George “Zhengyan” Kan for providing insightful comments and suggestions regarding lung cancer biology. We also thank the Computational Biology, Early Clinical Development, Tumor Cell Biology teams at the Pfizer Oncology Research Unit (ORD) for their support and feedback.

## References

[1] Ran FA, Hsu PD, Wright J, Agarwala V, Scott DA, Zhang F. Genome engineering using the CRISPR-Cas9 system. Nature Protocols. 2013;8(11):2281–308.

[2] Aguirre AJ, Meyers RM, Weir BA, Vazquez F, Zhang C-Z, Ben-David U, et al. Genomic Copy Number Dictates a Gene-Independent Cell Response to CRISPR/Cas9 Targeting. Cancer Discovery. 2016;6(8):914–29.

[3] Krijgsman O, Carvalho B, Meijer GA, Steenbergen RD, Ylstra B. Focal chromosomal copy number aberrations in cancer—Needles in a genome haystack. Biochimica et Biophysica Acta (BBA) - Molecular Cell Research. 2014;1843(11):2698–704.

[4] Munoz DM, Cassiani PJ, Li L, Billy E, Korn JM, Jones MD, et al. CRISPR Screens Provide a Comprehensive Assessment of Cancer Vulnerabilities but Generate False-Positive Hits for Highly Amplified Genomic Regions. Cancer Discovery. 2016;6(8):900–13.

[5] Meyers RM, Bryan JG, McFarland JM, Weir BA, Sizemore AE, Xu H, et al. Computational correction of copy number effect improves specificity of CRISPR–Cas9 essentiality screens in cancer cells. Nature Genetics. 2017;49(12):1779–84.

[6] Weck AD, Golji J, Jones MD, Korn JM, Billy E, Mcdonald ER, et al. Correction of copy number induced false positives in CRISPR screens. PLOS Computational Biology. 2018;14(7).

[7] Iorio F, Behan FM, Gonçalves E, Bhosle SG, Chen E, Shepherd R, et al. Unsupervised correction of gene-independent cell responses to CRISPR-Cas9 targeting. BMC Genomics. 2018;19(1).

[8] Gonçalves E, Behan FM, Louzada S, Arnol D, Stronach EA, Yang F, et al. Structural rearrangements generate cell-specific, gene-independent CRISPR-Cas9 loss of fitness effects. Genome Biology. 2019;20(1).

[9] Li T, Zhang Y, Patil P, Johnson WE. Overcoming the impacts of two-step batch effect correction on gene expression estimation and inference. 2021;

[10] Frank EH. Regression Modeling Strategies. Springer Series in Statistics. 2015;

[11] Pinheiro José C., Bates DM. Mixed-effects models in S and S-PLUS. Springer; 2000.

[12] Cullagh PM, Nelder JA. Generalized linear models. Chapman and Hall; 1999.

[13] Sasaki H, Hikosaka Y, Kawano O, Moriyama S, Yano M, Fujii Y. Evaluation of Kras Gene Mutation and Copy Number Gain in Non-small Cell Lung Cancer. Journal of Thoracic Oncology. 2011;6(1):15–20.

[14] Ferrer I, Zugazagoitia J, Herbertz S, John W, Paz-Ares L, Schmid-Bindert G. KRAS-Mutant non-small cell lung cancer: From biology to therapy. Lung Cancer. 2018;124:53–64.

[15] Goulet M-A, Cousineau D. The Power of Replicated Measures to Increase Statistical Power. Advances in Methods and Practices in Psychological Science. 2019;2(3):199–213.

[16] Zhu D, Li Y, Li H. Multivariate correlation estimator for inferring functional relationships from replicated genome-wide data. Bioinformatics. 2007;23(17):2298–305.

[17] Zhao J, Han Y, Li J, Chai R, Bai C. Prognostic value of KRAS/TP53/PIK3CA in non-small cell lung cancer. Oncology Letters. 2019;

[18] Palaniappan A, Ramar K, Ramalingam S. Computational identification of novel stage-specific biomarkers in colorectal cancer progression. PLoS ONE. 2016;11(5):e0156665.

[19] Boudhraa Z, Carmona E, Provencher D, Mes-Masson A. Ran GTPase: a key player in tumor progression and metastasis. Frontiers in Cell and Developmental Biology. 2020;

[20] Zohud BA, Guo P, et al. Importin 13 promotes NSCLC progression by mediating RFPL3 nuclear translocation and hTERT expression upregulation. Cell Death Dis. 2020; 11, 879.

[21] Hasan N, Ahuja N. The Emerging Roles of ATP-Dependent Chromatin Remodeling Complexes in Pancreatic Cancer. Cancers. 2019;11,1859.

[22] Anders S, Huber W. Differential expression analysis for sequence count data. Nature Precedings. 2010.

[23] Dempster JM, Pacini C, Pantel S, Behan FM, Green T, Krill-Burger J, et al. Agreement between two large pan-cancer CRISPR-Cas9 gene dependency datasets. 2019.

[24] Johnson WE, Li C, Rabinovic A. Adjusting batch effects in microarray expression data using empirical Bayes methods. Biostatistics. 2006;8(1):118–27.

[25] Leek JT, Johnson WE, Parker HS, Jaffe AE, Storey JD. The sva package for removing batch effects and other unwanted variation in high-throughput experiments. Bioinformatics. 2012;28(6):882–3.

[26] Gauthier J, Wu QV, Gooley TA. Cubic splines to model relationships between continuous variables and outcomes: a guide for clinicians. Bone Marrow Transplantation. 2019;55(4):675–80.

[27] Stone CJ, Koo CY. Additive splines in statistics. Proceedings of the American Statistical Association. Original pagination is p. 1985;45:48.

[28] Wu A, Xiao T, Fei T, Liu XS, Li W. Reducing False Positives in CRISPR/Cas9 Screens from Copy Number Variations. 2018;

[29] Lun A. Overcoming systematic errors caused by log-transformation of normalized single-cell RNA sequencing data. 2018;

[30] Doench JG, Fusi N, Sullender M, Hegde M, Vaimberg EW, Donovan KF, et al. Optimized sgRNA design to maximize activity and minimize off-target effects of CRISPR-Cas9. Nature Biotechnology. 2016;34(2):184–91.

[31] Wang T, Birsoy K, Hughes NW, Krupczak KM, Post Y, Wei JJ, et al. Identification and characterization of essential genes in the human genome. Science. 2015;350(6264):1096–101.

