## Supplementary Materials for "Computational correction of cell-specific gene-independent effects in CRISPR-Cas9 essentiality screens: REStricted CUbic SplinEs with Mixed Models (RESCUE-MM)"

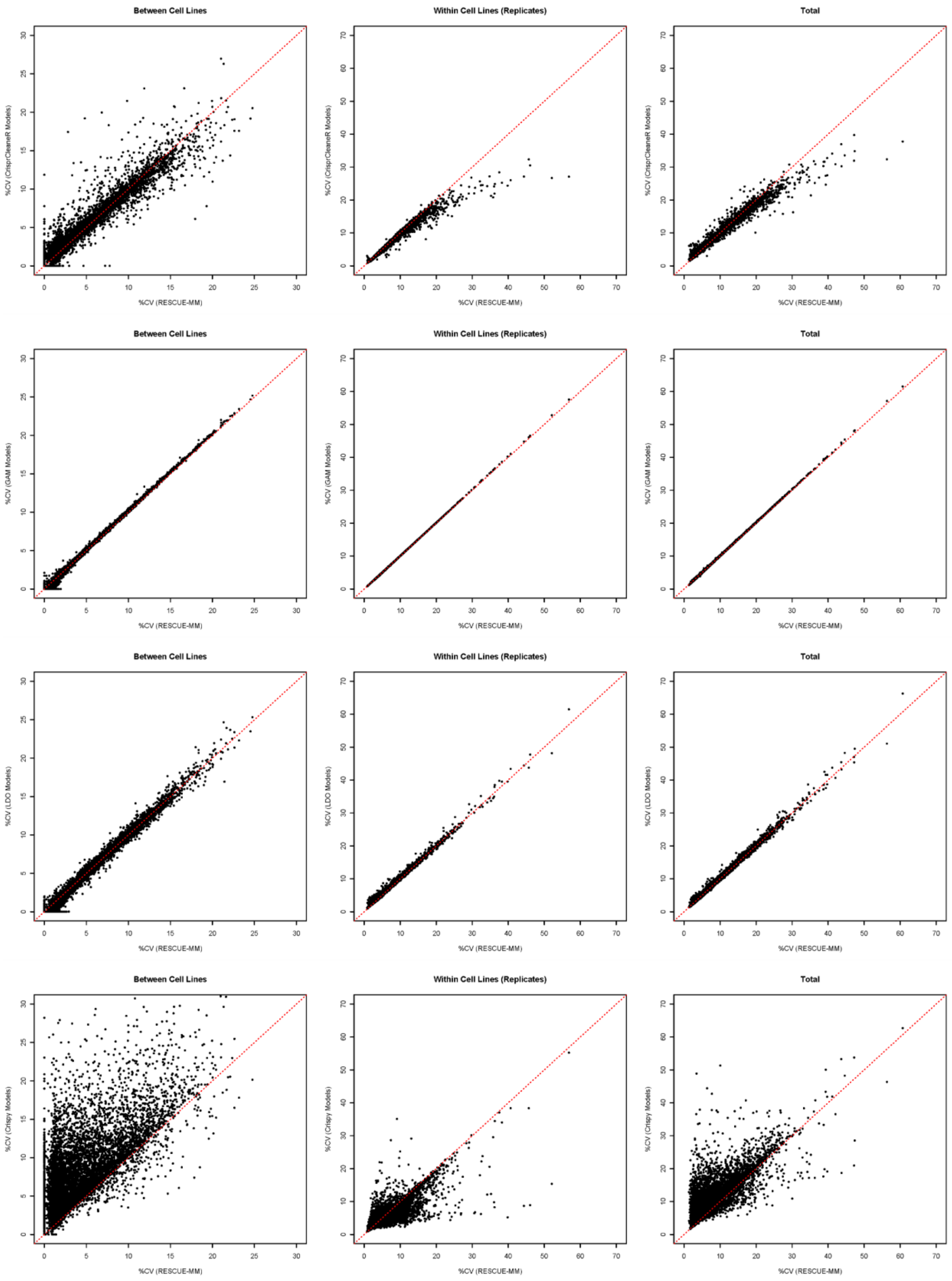


Suppl. Fig 1. %CV comparison for control guides, all methods (except CERES) versus RESCUE-MM.

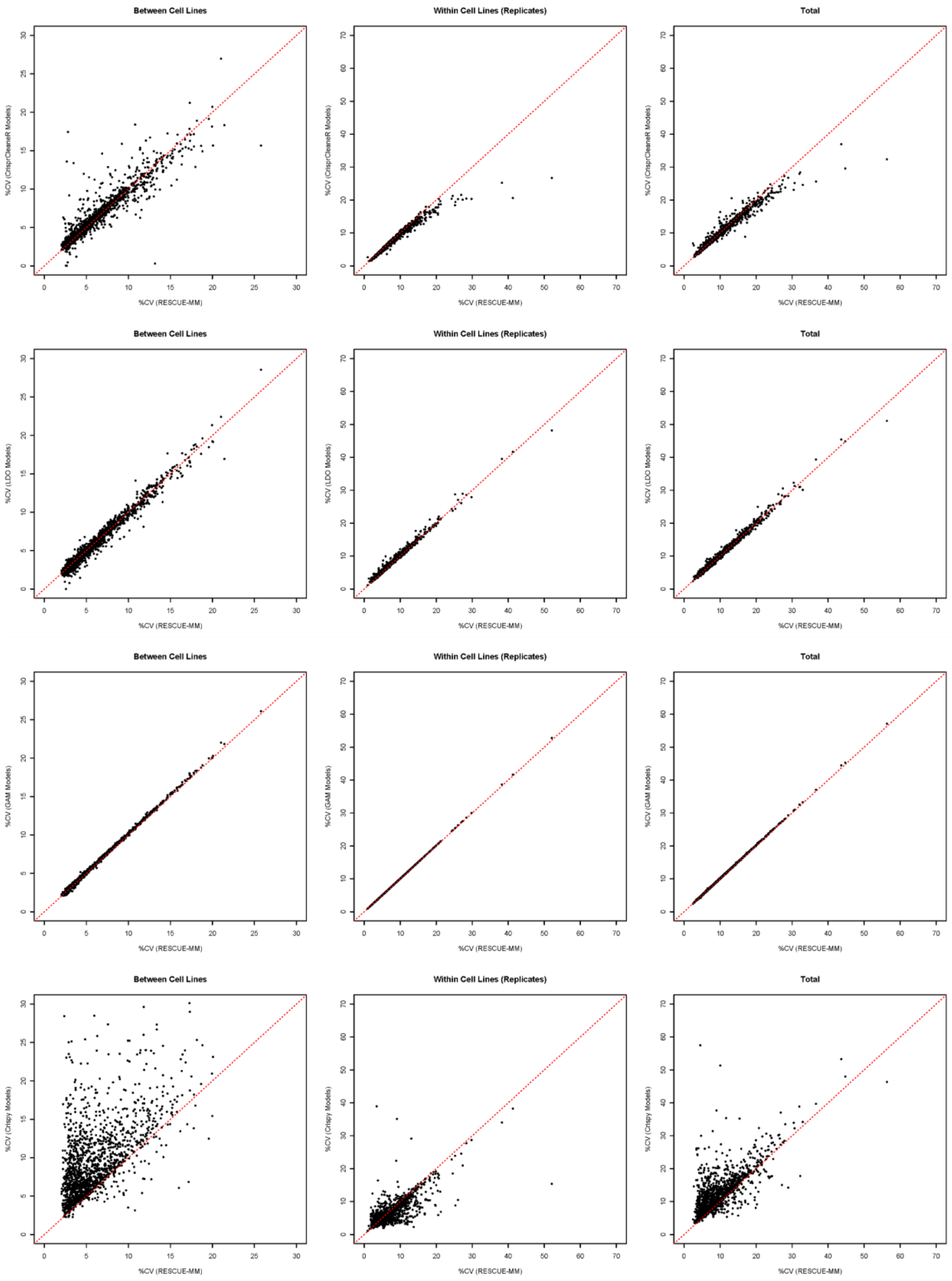


Suppl. Fig 2. %CV comparison for copy number affected guides, all methods (except CERES) versus RESCUE-MM.

| KRAS \ LMNTD1 | ACAATCCTGTGCAAACCGAA | | ATAGCTGAAGTGAATGTCAA | | GGATGAATCACCCATGATTG | | TTACGATGTTTGGAAGGAAT | |
|---|---|---|---|---|---|---|---|---|
| | **Unadjusted** | **Adjusted** | **Unadjusted** | **Adjusted** | **Unadjusted** | **Adjusted** | **Unadjusted** | **Adjusted** |
| AGAGGAGTACAGTGCAATGA | 0.277 | 0.139 | 0.305 | 0.131 | 0.301 | 0.172 | 0.444 | 0.308 |
| CGAATATGATCCAACAATAG | 0.294 | 0.143 | 0.231 | 0.022 | 0.399 | 0.274 | 0.439 | 0.282 |
| TCCCTTCTCAGGATTCCTAC | 0.341 | 0.201 | 0.364 | 0.184 | 0.370 | 0.243 | 0.496 | 0.356 |

Suppl. Table 1: Pearson correlation between targeted guides for KRAS and LMNTD1, neighbouring genes in the KRAS region, in unadjusted and copy number adjusted LFC using RESCUE-MM.

| sgRNA | Gene (Entrez-ID) | Chr | Loc | Str | RESCUE-MM | | RESCUE-GLMM | |
|---|---|---|---|---|---|---|---|---|
| | | | | | P-value | Adj. P-value | P-value | Adj. P-value |
| GCTCAACGACTCCATTGCCG | POLR3H (171568) | 22 | 41544022 | - | 9.43E-05 | 3.66E-01 | 4.30E-10 | 2.93E-05 |
| CGTAGCAAAGATCAACCGAG | PMPCA (23203) | 9 | 136417020 | + | 3.26E-07 | 2.22E-02 | 1.19E-08 | 4.05E-04 |
| GCGAAGATGGCGGAGAACAG | PFDN2 (5202) | 1 | 161118015 | - | 9.66E-06 | 1.11E-01 | 3.82E-08 | 8.67E-04 |
| GCCACCAAAACCCTCAACAG | SNU13 (4809) | 22 | 41675179 | - | 1.08E-03 | 1.00E+00 | 4.05E-07 | 6.89E-03 |
| ACCCTGGCCACGGTACGCCA | SNRPE (6635) | 1 | 203861663 | - | 7.17E-04 | 8.28E-01 | 1.03E-06 | 9.94E-03 |
| CCTTTGATAGATCTCTGCCG | TARS (6897) | 5 | 33456172 | + | 6.19E-04 | 7.74E-01 | 1.17E-06 | 9.94E-03 |
| TGCAGGCCGAGTACACCGAG | POLR2L (5441) | 11 | 842418 | - | 1.00E-04 | 3.66E-01 | 9.29E-07 | 9.94E-03 |
| TTCACTTGTGGCAAGATCGT | POLR2L (5441) | 11 | 842470 | - | 3.73E-05 | 1.96E-01 | 1.10E-06 | 9.94E-03 |
| CCTGGAGATGAGCAAGACCG | RUVBL2 (10856) | 19 | 49007098 | + | 2.85E-05 | 1.92E-01 | 1.36E-06 | 1.03E-02 |
| TTGCTGATGGAATTCAAGAA | PSMD11 (5717) | 17 | 32454529 | - | 1.84E-03 | 1.00E+00 | 1.54E-06 | 1.05E-02 |
| AGCTGGGTGTGAAGTCCCCA | TBL3 (10607) | 16 | 1975404 | + | 1.04E-03 | 1.00E+00 | 2.61E-06 | 1.37E-02 |
| CTGCAGGCCGAGTACACCGA | POLR2L (5441) | 11 | 842419 | - | 6.08E-04 | 7.74E-01 | 2.50E-06 | 1.37E-02 |
| TATATGTGATGATCTTAGAG | KARS (3735) | 16 | 75635731 | + | 1.08E-03 | 1.00E+00 | 2.53E-06 | 1.37E-02 |
| GCGGGGTCTCCGGAACCAAA | RPL35A (6165) | 3 | 197951208 | + | 8.86E-03 | 1.00E+00 | 3.09E-06 | 1.50E-02 |
| GCAGGAAGAAGAGAGCGAAG | VPS41 (27072) | 7 | 38869240 | - | 1.06E-03 | 1.00E+00 | 3.57E-06 | 1.62E-02 |
| ACAGCCAGGAGAAGGCCAAG | TSR2 (90121) | X | 54440737 | + | 3.00E-03 | 1.00E+00 | 4.93E-06 | 1.63E-02 |
| ACTTGAGGCAGAACTCGCAC | KAT5 (10524) | 11 | 65714691 | - | 1.78E-03 | 1.00E+00 | 5.02E-06 | 1.63E-02 |
| AGTGCTGGTAGAGCACCAGG | CDC45 (8318) | 22 | 19507394 | - | 2.88E-04 | 5.14E-01 | 4.13E-06 | 1.63E-02 |
| GGTTACACAGACATACCCGT | ATP6V1A (523) | 3 | 113784437 | - | 5.46E-04 | 7.59E-01 | 4.79E-06 | 1.63E-02 |
| TCAGGCCAAGAAGTACGCCA | PUF60 (22827) | 8 | 143821673 | - | 4.53E-05 | 2.20E-01 | 4.04E-06 | 1.63E-02 |
| TTTGCCTTCCCAGTACGTGG | TARS (6897) | 5 | 33457266 | - | 6.18E-05 | 2.63E-01 | 4.50E-06 | 1.63E-02 |
| GCAACCCACACACTTCGGAG | SRSF3 (6428) | 6 | 36596870 | - | 1.24E-04 | 3.83E-01 | 6.33E-06 | 1.96E-02 |
| CGTACGGGACACGATCGCCA | POP5 (51367) | 12 | 120581165 | - | 1.79E-04 | 4.35E-01 | 7.26E-06 | 2.15E-02 |
| ACGCAAAGCTGTCATCGTGA | RPL27 (6155) | 17 | 42998826 | + | 4.37E-04 | 6.70E-01 | 8.98E-06 | 2.55E-02 |
| AGTTCGATATTTCTCTGCGT | WEE1 (7465) | 11 | 9577215 | + | 2.77E-04 | 5.14E-01 | 1.03E-05 | 2.80E-02 |
| GCTGAACATGGTCTACCAGG | SNAPC4 (6621) | 9 | 136395667 | - | 2.95E-04 | 5.14E-01 | 1.26E-05 | 3.29E-02 |
| GCGTCTGGAAGGCAACACAG | RANGAP1 (5905) | 22 | 41274661 | - | 4.14E-06 | 7.04E-02 | 1.32E-05 | 3.34E-02 |
| GGTATTCGCGAATGGCCCCA | DYNC1H1 (1778) | 14 | 101986021 | - | 1.78E-04 | 4.35E-01 | 1.41E-05 | 3.44E-02 |
| CCAGCGCGCTACTTACAGTG | RPS13 (6207) | 11 | 17077429 | + | 6.01E-03 | 1.00E+00 | 1.59E-05 | 3.54E-02 |
| CCTGGAGTCCAGTATCACAG | DYNC1H1 (1778) | 14 | 102010846 | + | 2.09E-06 | 4.75E-02 | 1.61E-05 | 3.54E-02 |
| GAAGCAGCAGAAGAAGGAGG | UBE2M (9040) | 19 | 58558344 | - | 4.22E-04 | 6.68E-01 | 1.59E-05 | 3.54E-02 |

| | | | | | | | | |
|---|---|---|---|---|---|---|---|---|
| TTGATGGGTCAAAGTCCGGC | YJU2 (55702) | 19 | 4249241 | - | 8.24E-04 | 9.21E-01 | 2.02E-05 | 4.30E-02 |
| GCTGGCGGCTGCTCTCCTGG | SMC1A (8243) | X | 53409131 | + | 2.88E-03 | 1.00E+00 | 2.13E-05 | 4.40E-02 |
| GATGAAGTGCCCCTTGGACA | ORC6 (23594) | 16 | 46691115 | + | 2.74E-04 | 5.14E-01 | 2.35E-05 | 4.70E-02 |
| TGAGGACGTGAAGCGCACAG | RPN1 (6184) | 3 | 128650685 | - | 2.08E-03 | 1.00E+00 | 2.48E-05 | 4.83E-02 |
| TCTCACAGCCTGATGCCCAG | IPO13 (9670) | 1 | 43950148 | + | 1.05E-06 | 3.58E-02 | 8.87E-05 | 1.04E-01 |

Suppl. Table 2: Synthetic lethal sgRNAs in presence of KRAS mutation in lung cancer cell lines identified by RESCUE-MM and RESCUE-GLMM approaches. Benjamini-Hochberg p-value adjustment method was used.
